# Modality-independent effect of gravity in shaping the internal representation of 3D space

**DOI:** 10.1101/2020.03.23.003061

**Authors:** Morfoisse Theo, Herrera Altamira Gabriela, Angelini Leonardo, Clément Gilles, Beraneck Mathieu, McIntyre Joseph, Tagliabue Michele

## Abstract

Human 3D perception of visual objects is flawed by distortions, which are influenced by non-visual factors, such as gravitational vestibular signals. Whether gravity acts specifically on the visual system or at a higher, modality-independent, level of information processing remains unknown. To test these modality-specific vs modality-independent hypotheses, we performed experiments comparing visual versus haptic 3D shape perception in normo-gravity and microgravity. The results obtained for upright and supine posture in 1g show that visual and haptic perceptual anisotropies are systematically in opposing ego-centered, but not gravity-centered, directions suggesting they share a common origin. On the other hand, microgravity significantly modulates both visual and haptic perceptual distortion in the same direction. Overall, our results show a clear link between the visual and haptic perceptual distortions and demonstrate a role of gravity-related signals on a modality-independent internal representation of 3D space used to interpret incoming sensory inputs.

## Introduction

Perception of tridimensional (3D) objects includes the ability to determine an item’s location in space, as well as its geometrical properties, such as the relative size along each of three dimensions and the relative orientation of its edges. Given its importance for interacting with the physical world, 3D object perception has been deeply investigated. Visual perception has received the most attention, showing how various features of the stimuli such as disparities, size, occlusions, perspective, motion, shadows, shading, texture and blur all influence 3D visual perception (Welchman 2016) and how *internal models* shape the interpretation of the sensory signals (Curry 1972, Kersten and Yuille 2003, Kersten et al. 2004, Lee 2015).

Despite its critical importance to perception and action, visual perception suffers from measurable distortions: height underestimation with respect to width, also known as the horizontal-vertical, or “L”, illusion (Avery and Day 1969) and a systematic depth un-derestimation (Loomis and Philbeck 1999, Todd and Norman 2003). Non-visual factors, such as gravity, also appear to affect visual perception. For example, tilting the body with respect to gravity affects objects recognition (Leone 1998, Barnett-Cowan et al. 2015), orientation and distance perception (Marendaz et al. 1993, Harris and Mander 2014), and other phenomena such as the tilted frame illusion (Goodenough et al. 1981, Howard 1982), the oblique effect (Lipshits and McIntyre 1999, Luyat and Gentaz 2002, McIntyre and Lipshits 2008) and some geometric illusions (Prinzmetal and Beck 2001, Clément and Eckardt 2005). Furthermore, weightlessness significantly alters the perception of stimulus size and shape, especially in tasks involving depth, during both short-term (Villard et al. 2005, Clément and Bukley 2008, Clément et al. 2008, Harris et al. 2010, Clément and Demel 2012, Clément et al. 2016, Bourrelly et al. 2016) and long-term (Clément et al. 2012-2013, De Saedeleer et al. 2013, Bourrelly et al. 2015) exposure.

One hypothesis to explain gravity-related changes in visual perception is that gravity affects both the eye movements underlying visual exploration (Clément et al. 1986, Reschke et al. 2017-2018) and eye positioning that contribute to the estimation of the visual eye-height, a key reference within the visual scene (Goltz et al. 1997, Bourrelly et al. 2016). Gravity’s influence on oculomotor control should specifically affect visual perception, although weightlessness might also induce distinct distortions in other sensory modalities. An alternative hypothesis is that gravity does not affect visual signals *per se,* but rather affects an internal representation of space (Clément et al. 2009, 2012), based on prior knowledge, that serves to interpret those signals, independent of the sensory system from which they come (Wolbers et al. 2011, Loomis et al. 2013). An example, among many, of the use of an internal model of space for perception is the famous ‘Ames room’ illusion, where the persons’ size is misperceived due to the use of the inappropriate prior that the room is rectangular (O’Reilly et al. 2012). A direct implication of this second hypothesis is that microgravity should distort all spatial perceptions in the same way, regardless of the sensory modality. Because previous studies in microgravity were focused on visual tasks only, however, these proposed hypotheses have never been tested.

To investigate these two assumptions, we first compared distortions of visual versus haptic perception of 3D shape in a normal, upright posture on Earth. Next, we studied the effect of changing the subject’s orientation with respect to gravity to assess whether any visual or haptic distortions are egocentric or gravity-centric. Third, we tested the consequences of removing the effects of gravity by performing both haptic and visual experiments in weightlessness during parabolic flight.

## Materials and Methods

In an analogy with previous experiments on visual perception (Clément et al. 2008, 2013), our paradigm was conceptually designed to detect distortions in the perception of a purported 3D cube. The sequential nature of haptic perception induced us, however, to focus each trial on the comparison of the relative size between two out of three possible dimensions. In both the visual and the haptic cases, the task consisted of adjusting one side of the rectangle to match the other, to form a square. The adjustments were performed using a trackball hold in the left hand. Subjects pressed a button on a trackball when they perceived the object to be perfectly square.

For the haptic tasks, subjects were asked to close their eyes and to feel, through haptic sense only, a rectangular cutout in a rigid, virtual plank generated by a Force Dimension Omega.3 haptic robot (Figure 1A). This manipulandum was able to simulate the presence of a 3D object by applying the appropriate contact forces on the right hand of the subject when he/she performed exploration movements aimed at perceiving its shape and size. During each trial the robot constrained the subject’s hand movement to lie within the plane of the virtual plank and to remain inside the rectangle prescribed by the virtual cutout. To allow direct comparisons between the experimental results from haptic and visual tests, an analogous bi-dimensional task was also used for visual perception. Subjects were shown planar rectangles with different orientations in 3D space, without being able to manually explore it. For trials involving visual perception, an Oculus Rift virtual reality headset was used to provide a stereoscopic view of the virtual object. The visual environment was dark and the shapes were represented by light-gray frames. For both sensory conditions, the virtual object was located approximatively 40 cm in front of the subject’s right shoulder.

**Figure 1:**
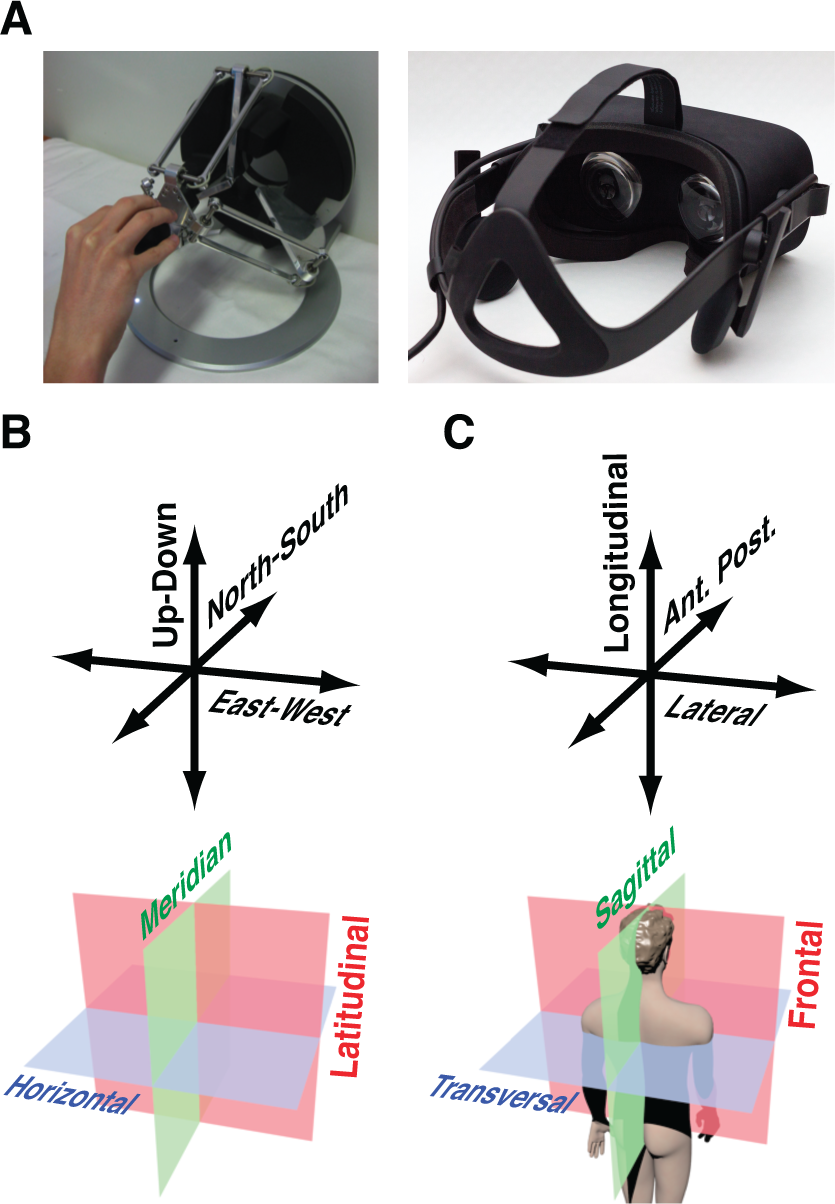
A) Haptic device and virtual reality headset used for the haptic and visual experiments, respectively. In panels B and C are reported the name of the orthogonal directions defined inan external, gravity-centered (Up-Down, UD; East-West, EW; North-South, NS) and egocentric, body-centered (Longitudinal, Lateral, and Anterior-Posterior) reference frames respectively. The bottom part of the figure represents the planes in which the task is performed expressed in the gravity-centered (Horizontal, Meridian and Latitudinal) and body-centered (Transversal, Sagittal and Frontal) reference frames.

Although there were no instructions to work quickly, subjects were asked to attempt to perform each trial in a fixed time window (20 s for all experiments except those performed on board the parabolic flight plane, for which a 10 s time window was used). An audible cue indicated to the subject when the end of the allotted time was approaching. The apparatus recorded the subject’s final responses (dimensions of each rectangle judged to be square), which is the main output of the tests. For the haptic tasks, the movements of the subject’s hand and the contact forces applied against the virtual constraints were also recorded via the haptic device.

The use of bi-dimensional tasks allowed the estimation of the perceptive distortion in one plane at a time. Subjects in our experiments judged the squareness of rectangles lying in each of three anatomical planes: frontal, sagittal, or transversal (see bottom part of Figure 1C). The combination of the three possible planes and the two rectangle dimensions resulted in six different geometric configurations that the subject had to deal with. They are represented in the upper part of Figure 2. At the beginning of each trial, an audio command told the subject in which anatomical plane the rectangle was lying and which of the two dimensions of the rectangle had to be adjusted. In our paradigm, the reference dimension was always 40 mm, but subjects were not aware of this fact. The initial length of the adjustable side was randomly selected between 15, 25, 35, 45, 55, and 65 mm. Subjects performed five series of trials in all; each series being composed of a random permutation of the six geometric configurations (total number of trials per condition: 30).

**Figure 2:**
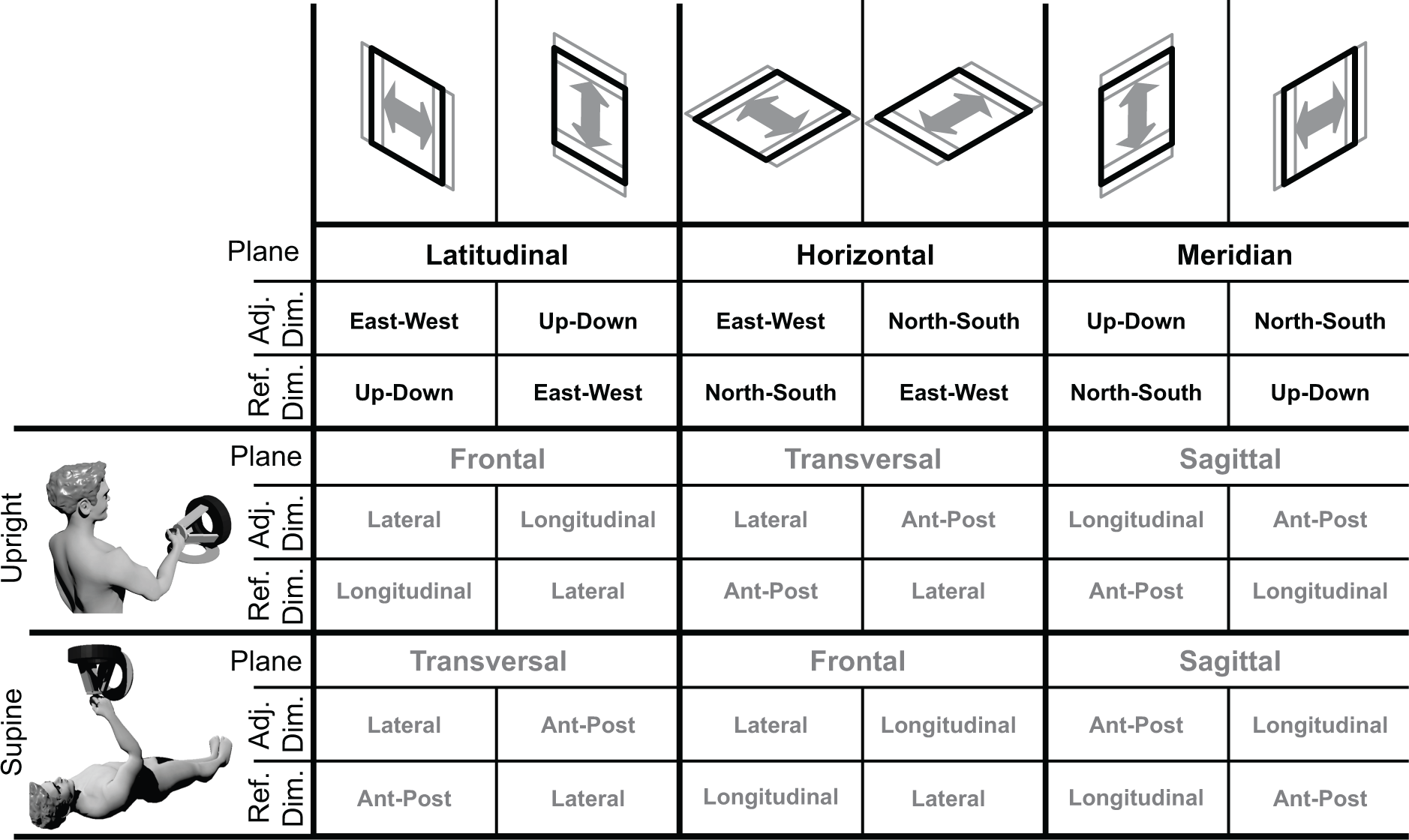
Geometrical Configurations. The first row represents the six geometric configurations, which correspond to the combination of the three planes in which the rectangle could lie and the two different dimensions of the rectangle that the subject had to adjust. The black bold text corresponds to the definitions, in the gravity-centered reference frame, of the task planes, as well as of the adjustable and reference dimensions of each rectangle. Those definitions are independent of the postural condition of Experiment 2 (Upright and Supine) represented in the lower part of the table. For each combination of geometric and postural conditions, the table reports with gray bold text the anatomical plane in which the task is performed as well as the anatomical direction of the adjustable and reference dimensions of the rectangles.

### Experiment 1: Effect of Sensory Modality

To study the differences and similarities between haptic and visual perception of 3D shapes in normo-gravity, 18 seated subjects (8 males, 10 females, aged 29±9) performed the task for all six geometrical configurations in each of the two sensory conditions: Haptic and Visual. The order of the two sensory conditions was randomized across subjects.

### Experiment 2: Effect of Body Orientation

To study the perceptive distortions of both haptic and visual senses and whether the information is encoded in an ego-centric (body-centered) or exo-centric (gravity-centered) reference frame, a group of 18 subjects (9 males and 9 females, aged 25.5±5 years) performed the haptic task while seated (Upright) and while lying on the back (Supine), while a second group of 18 subjects (11 male and 7 female, aged 24±4 years) performed the visual task in the same two postures (Upright and Supine). For the Supine posture, subjects laid on a medical bed. The two postures are represented in Figure 2 together with the respective correspondence between exo-centric and ego-centric references. The virtual object was placed always at the same distance from the subject’s shoulder, independent of the posture. In order to compensate for possible learning effects, the order of the postural conditions was randomized in both sensory conditions.

### Experiment 3: Effect of Weightlessness

To study the role of gravitational cues in the encoding of haptic and visual signals we performed the haptic (18 subjects: 10 males, 8 females, aged 38±11 years) and visual (18 subjects: 9 males, 9 females, aged 41±11 years) paradigm in normal gravity (1G) and during the weightlessness phases of parabolic flight (0G). For the haptic experiment, a third condition was added: the subjects were also tested in normal gravity, but with the arm supported by a strap (Supp.), to differentiate the biomechanical effect of gravity on the arm from the gravitational stimulation of graviceptors, such as the otoliths.

Parabolic flight provides short intervals (∼20s) of weightlessness within a stable visual environment inside the airplane, bracketed by periods of hyper-gravity (1.6 - 1.8 G) just before and just after each period of weightlessness. Given the short duration of 0G phases during parabolic flight, the subjects were trained to perform the task in about 10 seconds (two tasks per parabola). Since each subject performed the experiment during 15 consecutive parabolas, he or she could perform all 30 trials per condition.

All experimental conditions were performed inflight onboard the Novespace Zero-G airplane in order to minimize possible undesired changes in uncontrolled factors. The 1G and Support conditions were tested during the level-flight phase just preceding the first parabola or just following the last parabola of its session, depending on the subject. The subjects were very firmly restrained with belts so that their relative position with respect to the apparatus and the virtual rectangles did not vary between gravitational conditions.

### Data analysis

For each trial, *t*, the error, *ε*, between the length of the adjustable and reference sides of the rectangle was computed. If the exo-centered definition of the three dimensions (*EW*, *NS,* and *UD*) of Figure 1B is used, the errors of the six geometric configurations are defined as *EW-UD*, *UD-EW*, *EW-NS*, *NS-EW*, *UD-NS,* and *NS-UD*, where the minuend and the subtrahend are the adjustable and reference dimensions respectively.

Table *1* shows how the perceptive distortion associated with each of the three dimensions contributes to the error made on the six geometric configurations. Positive errors correspond to underestimations of the adjustable dimension and/or to overestimations of the reference dimension. Thus, the present experimental paradigm, similar to the one previously used by Clément et al. (2008, 2013), allows the quantification of the perceptive distortions of one dimension relative to another, but cannot lead to a measure of the absolute perceptive distortions for each dimension separately. Nevertheless, we have devised a methodology that allows one to visualize the 3D patterns of distorsion as an elongated box compared to an ideal cube (see Appendix I).

**Table 1:** Definition of the squaring errors for all six geometrical configurations of the task.

| Plane | Adjustable dimension | Reference dimension | Task error |
| --- | --- | --- | --- |
| Latitudinal | EW | UD | $\varepsilon_{EW-UD} = \varepsilon_{EW} - \varepsilon_{UD}$ |
| | UD | EW | $\varepsilon_{UD-EW} = \varepsilon_{UD} - \varepsilon_{EW}$ |
| Horizontal | EW | NS | $\varepsilon_{EW-NS} = \varepsilon_{EW} - \varepsilon_{NS}$ |
| | NS | EW | $\varepsilon_{NS-EW} = \varepsilon_{NS} - \varepsilon_{EW}$ |
| Meridian | UD | NS | $\varepsilon_{UD-NS} = \varepsilon_{UD} - \varepsilon_{NS}$ |
| | NS | UD | $\varepsilon_{NS-UD} = \varepsilon_{NS} - \varepsilon_{UD}$ |

#### Estimation of 3D perceptual distortion

Table *1* shows that the error in estimating one dimension has opposite effects for the two tasks performed within a given plane. For instance, an overestimation of the NS dimension should result in negative and positive errors in the NS-EW and EW-NS tasks, respectively. These relationships appear to be confirmed by the experimental results (Figure 4a), because this hypothesis accounts for 96% of the data variance. It follows that the theoretical relationships below are valid:

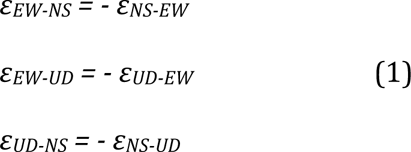

Exploiting this property, it was possible to combine the five errors obtained for one geometric condition, with the additive inverse of the five errors obtained for the other geometric condition performed in the same plane. This allowed computing the combined mean and the variance of the errors for each of the three planes (Horizontal, Latitudinal, and Meridian), instead of individually for each of the 6 geometrical configurations of the task. This technique has the considerable advantage of being more robust, because it is based on 10 samples instead of only 5.

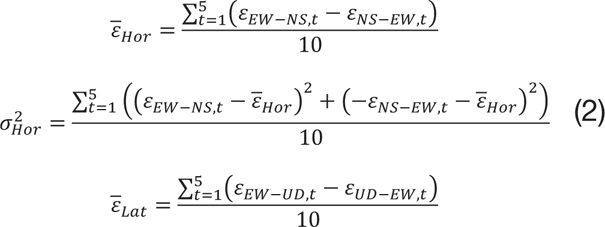

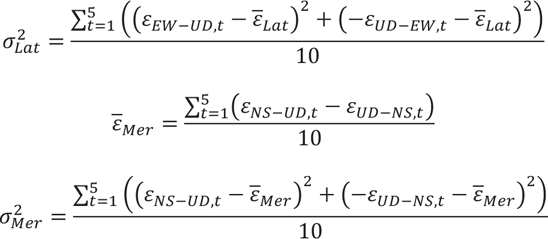

With the above formulas, one can characterize perceptual distortions in each of the three different planes as illustrated in Figure 3. By our convention, a rectangle lying in one of the two vertical planes (Meridian or Latitudinal) is associated with a positive error (stubby rectangle) if the vertical dimension is smaller than the other dimension. In the horizontal plane, a positive error (stubby rectangle) corresponds to the depth (NS dimension) being smaller than the width (EW dimension). It is worth noting that if the subject produced a “stubby” rectangle (positive errors) this means that he/she perceived a square to be “slender”, and vice versa. The global variance was computed as the average of the three planar variances.

**Figure 3:**
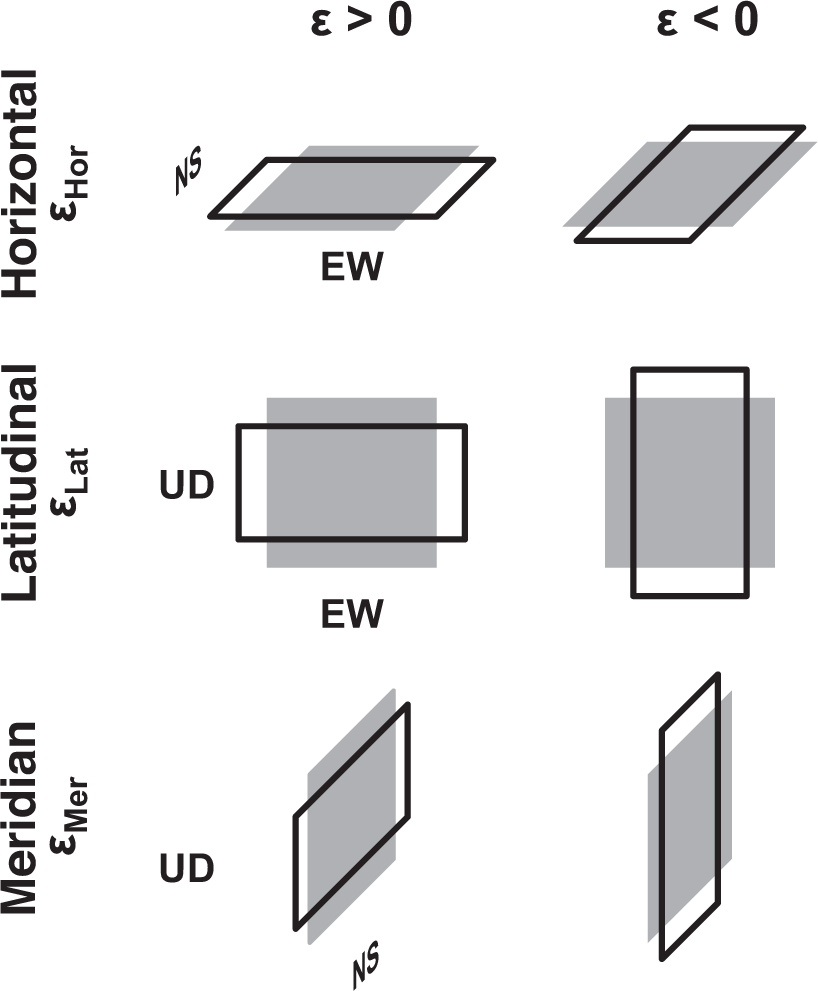
Sign conventions for the errors in the Horizontal, Latitudinal and Meridian planes. The gray squares represent the correct answer (i.e. a square). The black lines represent the distorted answers. Positive planar error values correspond to “stubby” rectangles. Negative values correspond to “slender” rectangles. The same conventions are used for the error expressed in the body-centered planes. In this case, Anterior-Posterior, Lateral and Longitudinal directions replace NS, EW and UD, respectively. Transversal, Frontal and Sagittal replace Horizontal, Latitudinal and Meridian planes, respectively.

The estimation of the three planar errors is then improved by considering that if the (distorted) metrics used to compare distances in 3D space are locally smooth and consistent for the different dimensions in space, the three planar errors *ε* are not independent and that, given the sign conventions of Figure 3, they should fulfill the following relationship

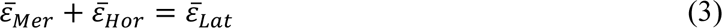

Note that equation 3 is a particular case of the formula describing a plane, a*x* + b*y* + c*z* = d, where a = b = 1, c = −1 and d = 0. Thus, if the metrics in each plane are consistent with each other, the vectors of measured planar errors 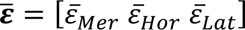 should fall on that plane and points outside the plane can be considered to be noise. By projecting the individual vectors 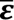 onto the plane corresponding to equation 3, this noise is effectively filtered out. Using the resulting 2D representation of the distortion is a conservative choice, especially when comparing their orientation in different conditions, because the 3D representation may lead to consider distortion directions and components of data variability that have no functional meaning. On average, the data projected on the plane of equation 3 account for 98% of the variance of the original data, suggesting that the recorded responses tend to well fulfill this constraint.

We used the same equations (1-3) to compute the analogous parameter in the ego-centric reference frame after having replaced the externally defined planes and directions with the body-centered planes (*Transversal*, *Frontal,* and *Sagittal*) and directions (*Lateral*, *Anterior-Posterior,* and *Longitudinal*) as shown in Figure 2. Table *2* shows the relationships between the planar distortions defined in the body-centered and gravity-centered reference frame for the Upright and Supine posture.

**Table 2:** Relationship between ego- and exo-centrically defined distortions for the Upright and Supine condition.

|  |  |  |  |
| --- | --- | --- | --- |
| **Upright** | $\bar{\varepsilon}_{Mer} = \bar{\varepsilon}_{Sag}$ | $\bar{\varepsilon}_{Lat} = \bar{\varepsilon}_{Fro}$ | $\bar{\varepsilon}_{Hor} = \bar{\varepsilon}_{Tra}$ |
| **Supine** | $\bar{\varepsilon}_{Mer} = -\bar{\varepsilon}_{Sag}$ | $\bar{\varepsilon}_{Lat} = \bar{\varepsilon}_{Tra}$ | $\bar{\varepsilon}_{Hor} = \bar{\varepsilon}_{Fro}$ |

#### Distortion direction

The 2D vector resulting from the projection of 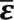 to the plane of equation 3 was computed for each subject. To estimate the mean direction of the distortion in each experimental condition, the vectorial average of the 2D vectors was computed.

#### Reaction forces during haptic task

To estimate changes of the contact forces between gravitational conditions in the haptic tasks, we computed the average of the reaction forces generated by the haptic device when the subject’s hand was in contact with the edges of the virtual cutout or when the hand tried to move out of the task plane.

#### Microgravity effect and modeling

To quantify the effect of microgravity on the perceptive distortions, for each subject the mean planar error in 1G was subtracted from the corresponding error in 0G:

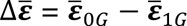

computed separately for the visual and the haptic modalities. As for the mean planar error, a 2D vector representation was computed projecting each individual 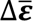 onto the plane of equation 3. To predict the perceptive distortion expected in microgravity under the hypothesis that the 0G effect was identical for the haptic and visual modalities, we averaged all 2D vectors representing the measured individual microgravity effects from both the haptic and visual experiments, combined. The obtained vector was then added to the individual visual and haptic distortions measured in normo-gravity conditions. We then compared these predictions to the distortions measured in 0G for the visual and haptic modalities.

#### Statistical analysis

For each experiment, we first tested the significance of the distortions by testing for each plane whether the constant errors were on average different from zero (two-sided Student’s t-test). Then, we performed repeated-measures ANOVA on the constant and variable errors. The sign conventions (Figure 3) being arbitrary, they allow a rigorous comparison of the perceptive distortion within a given plane, but they do not allow the comparison between different planes. For this reason, in the statistical analyses, the results on each plane were tested with independent ANOVAs.

<u>Experiment 1:</u> For each of the 3 task planes we tested for an effect of Sensory Modality on the perceptive distortion as a single within-subject independent factor with two levels (Haptic, Visual).

<u>Experiment 2:</u> We tested for an effect of Body Posture as a within-subject independent factor with two levels (*Upright*, *Supine*) in separate ANOVAs for each group/sensory modality (Visual and Haptic). Note that this separation is justified by the hypotheses being tested, for which cross effects between posture and modality would have little meaning. To test whether distortions are tied to an ego-centric or gravity-centric reference frame, we defined the task planes both in terms of anatomical axes and world axes. Invariance of distortion (lack of a statistical difference) for the anatomically defined planes, but not the world-defined frames, would indicate that the distortions are primarily egocentrically rather than exocentrically aligned and vice-versa.

<u>Experiment 3:</u> For each of the 3 task planes we tested for an effect of Gravity on shape perception as a single within-subject independent factor with three (1G, 0G, Supported) and two (1G, 0G) levels for the haptic and visual experiment respectively.

Before performing each ANOVA, we tested for normality and homogeneity of the distributions using the Kolmogorov-Smirnov and Levenes tests, respectively. To achieve the normal distribution for the response variability, the standard deviations were transformed by the log(σ+1) function (Tagliabue and McIntyre 2011). For the errors expressed in both exocentric and ego-centric reference frames the data were distributed normally (all p>0.20) and the data variability was similar among all conditions (all p>0.50).

To test whether the direction of the distortion (2D vectorial average) differs between two conditions, a bootstrap technique was used: 10000 re-samplings were used to estimate the statistical distribution of the angle θ between the vectorial average of two conditions, and to compute the probability of error in rejecting the null hypothesis, H_0_, that θ=0. The same technique was used to compare the global amplitude of the distortions.

Following a Bayesian approach, taking into account a prior uniform distribution of all possible angles (θ range ±180°), we evaluated the ratio, R_0/1_, between the probability to obtain the observed data under the null hypothesis, H_0_, and the probability under the alternative hypothesis, H_1_, that θ≠0 (Wagenmakers et al. 2018).

## Results

### Experiment 1: Haptic and Visual Perception

Figure 4A shows that for the six geometric configurations of the squaring task (see methods) the subjects made systematic errors in both visual and haptic conditions. The comparison of the errors made using haptic information alone versus visual information alone shows consistent, opposing results for the two sensory modalities. Hence, in each task, when subjects made on average significant positive errors in the haptic condition, they made negative errors in the visual condition, and vice-versa. Figure 4B represents the more robust evaluation of the planar distortions obtained by considering the constraints existing between the errors performed in the six squaring tasks (see methods equation 1-3). The amplitude of the distortion was significantly different from zero for both visual and haptic perception in the Sagittal (visual: F_(17)_=5.86, p<10^−4^, haptic: F_(17)_=-8.10, p<10^−6^) and Transversal plane (visual: F_(17)_=-7.22, p<10^−5^, haptic: F_(17)_=9.22, p<10^−6^), but in the Frontal plane none of the modality was significantly different from zero (visual: F_(17)_=-1.26, p=0.22, haptic F_(17)_=-0.57, p=0.58). Sensory modality had a significant effect in the Sagittal (F_(1,17)_=60.8, p<10^−5^) and Transversal (F_(1,17)_=94.96, p<10^−6^) planes, but not in the Frontal plane (F_(1,17)_=0.14, p=0.71). Remarkably, the significant perceptive distortions in the Sagittal and Transversal planes were in opposite directions between the two sensory conditions. When using haptic sense, subjects produced rectangles with the Anterior-Posterior dimension smaller than the Longitudinal and Lateral dimension, while, when using vision, they made rectangles with the Anterior-Posterior dimension larger than the Longitudinal and Lateral dimension.

**Figure 4:**
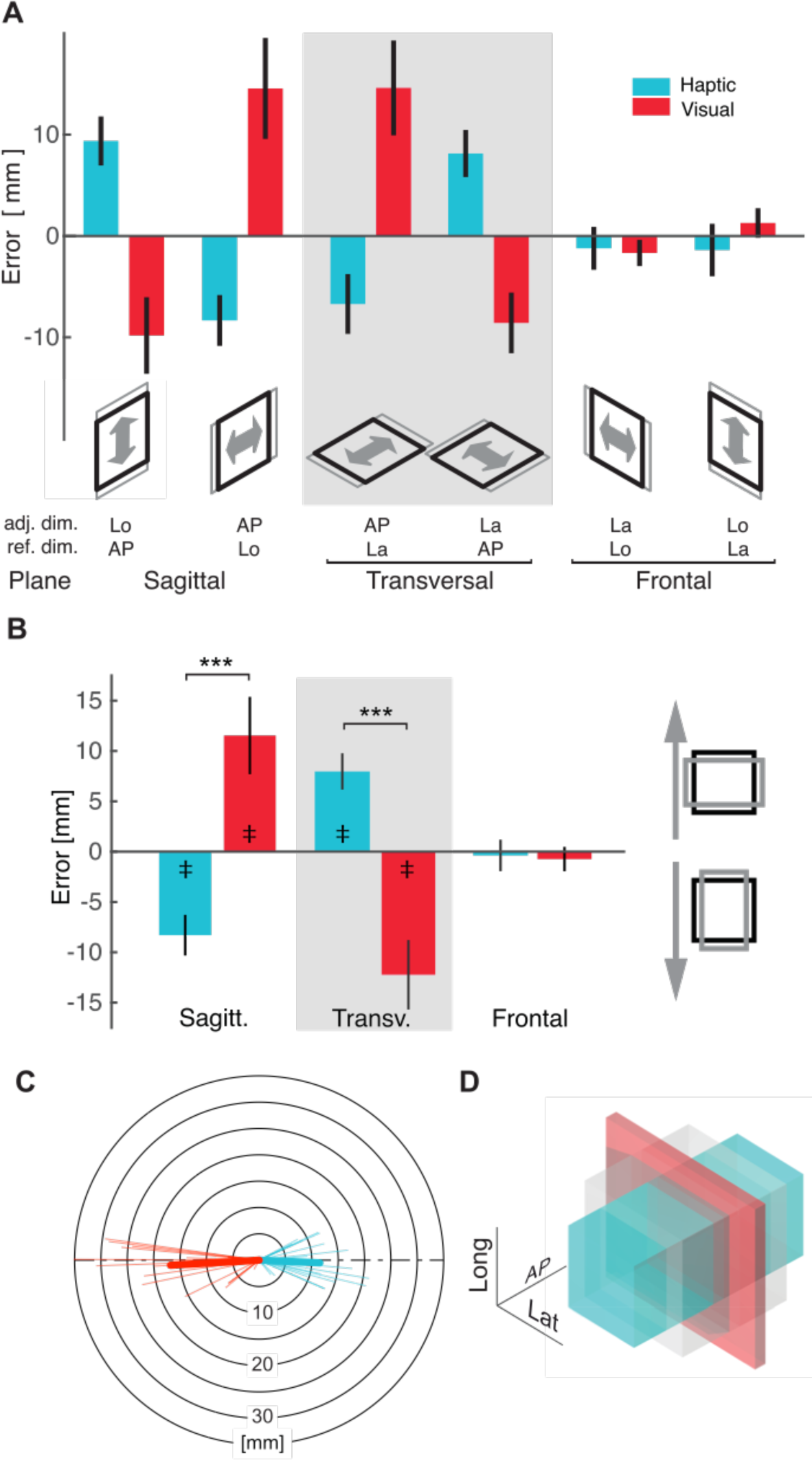
**A**) Errors for the task performed in each of the six geometrical conditions using haptic information only (light blue bars) or visual information only (red bars). Each geometrical condition is characterized by the plane in which the rectangle lies (sagittal, transversal, frontal), and by which direction within the plane was adjustable or held constant: Longitudinal (Lo), Anterior-Posterior (AP), and Lateral (La). Positive errors correspond to the final size of the adjustable dimension being greater than the reference dimension. Vertical wiskers represent 95% confidence intervals. A significant difference between the two tasks performed in the same plane is indicative of an important perceptive distortion in that specific plane. **B**) Distortions in the three task planes for haptic and visual conditions. *** : p<10^−3^ in the ANOVA testing the modality effect. ‡ : p<10^−3^ for the t-test ascertaining differences from zero. **C**) Vectorial representation of individual responses in the Sagitt+Transv=Frontal errors space. Thicker vectors represent the average vectorial response. Dashed line indicates the Sagitt+Transv=0 axis. **D**) Illustration of how a cube (gray shape) would be perceived by the subjects when using haptic or visual information alone, respectively. For illustration purposes, the distortions of this panel are scaled up by a factor of 5.

The vectorial representation of the individual errors in Figure 4C shows that the perceptive distortions for the two sensory modalities appear along the same axis, but in opposite directions: average angle θ between the visual and haptic distortion is 174±5° and not significantly different from 180° (bootstrap p=0.13). Taking into account all possible orientations for the two groups of vectors, the observed results are 14 times more likely under the hypothesis that distortions of the two senses are in opposite directions (H_0_: θ =180°), than under the alternative hypothesis (H_1_), i.e. θ≠ 180°. The amplitude of the distortion is larger for visual than for haptic sense (bootstrap p=0.02). The illustration of the perceptive distortion corresponding to the two sensory modalities is reported in Figure 4D and suggests that the haptic and visual perceptive distortions would mainly consist of a depth overestimation and underestimation for the haptic and visual sense, respectively. The method used to compute the parallelepiped dimensions is described, with its limitations, in Appendix 1.

If the amplitude of the perceptive biases (constant components of the errors reported in Figure 4 A-C) appear smaller for the haptic than for the visual modality, the latter is characterized by a clearly smaller intra-personal variability of the responses (α_hapt_=6.1±2.6 mm, α_vis_=4.2±2.2 mm, sensory modality effect: F_(1,17)_=12.02, p<10^−2^), corresponding to a higher precision for the visual than for the haptic task.

In summary, Experiment 1 shows clear differences in the patterns of visual and haptic distortions. Distortion appeared primarily in the sagittal and transversal planes and were in opposite direction for the two sensory modalities (contraction and expansion of perceived depth for visual and haptic, respectively).

### Experiment 2: Effect of Body Orientation

The responses of the subjects upright were characterized by constant errors similar to those observed in Experiment 1 (Experiment effect: Wilks’ Lambda=0.85, F_(4,32)_=1.35, p=0.27). The left columns of Table 3 and left panels of Figure 5 show that for both haptic and visual experiments the planar distortions appear consistent between postures if expressed ego-centrically: we observed no statistically significant effects of posture on the errors for any of the three planes when expressed in body-centered reference frame, nor on the distortion magnitude (boostrap for haptics p=0.76 and vision p=0.75) and direction (boostrap for haptics p=0.52 and vision p=0.36; R_0/1_=10.1 for haptics and R_0/1_=21.8 for vision). On the other hand, if the errors are represented in terms of exo-centrically defined planes, i.e. fixed with respect to gravity (last three columns of Table 3 and right panels of Figure 5), a clear effect of posture can be observed in all planes for both sensory modalities and on the distortion orientation (bootstrap p<10^−4^ for haptics and vision; R_0/1_<10^−4^ for haptics and vision).

**Figure 5:**
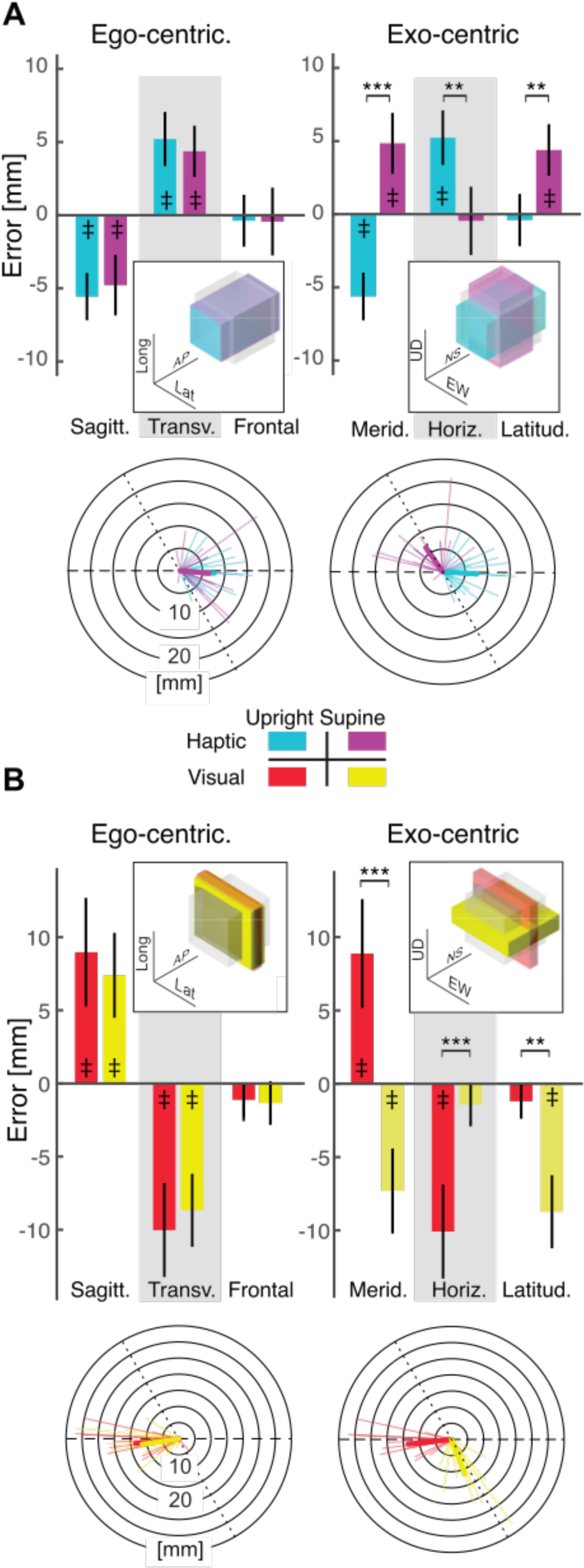
Errors within each plane when the subjects are seated normally (Upright) or lying Supine. The upper (**A**) and lower (**B**) panels represent the results for the Haptic and Visual modalities, respectively. The left panels represent the errors per anatomical, ego-centered plane. The right panels represent the data per exo-centered (fixed with respect to gravity) plane. ** : p<10^−2^ and *** : p<10^−3^ in the ANOVA. † and ‡ : p<10^−2^ and p<10^−3^ for the t-test ascertaining differences from zero. Vertical wiskers represent 95% confidence intervals. In each barplot the inset illustrates the corresponding 3D perceptive distortion (amplified x5) of a cube. The circular vectorial plots represent individual responses in the Sagitt+Transv=Frontal errors space. Thicker vectors represent the average vectorial response. Dashed diameters indicate the Sagitt+Transv=0 (or the Merid+Horiz=0) axis. Dotted diameters indicate the Saggitt=Frontal (or Merid=Latitud) axis, corresponding to a 90° rotation of the distortions pattern.

**Table 3:** Results of ANOVA for the posture effect on the planar perceptive distortion.

|  | Sagittal | Transversal | Frontal | Meridian | Horizontal | Latitudinal |
| --- | --- | --- | --- | --- | --- | --- |
| Haptic | F(1,17)=0.40<br>p=0.53 | F(1,17)=0.58<br>p=0.46 | F(1,17)=0.001<br>p=0.97 | F(1,17)=52.28<br>p<10<sup>-5</sup> | F(1,17)=13.01<br>p=0.002 | F(1,17)=12.18<br>p=0.003 |
| Visual | F(1,17)=2.00<br>p=0.18 | F(1,17)=1.32<br>p=0.27 | F(1,17)=0.15<br>p=0.70 | F(1,17)=25.46<br>p<10<sup>-3</sup> | F(1,17)=19.92<br>p<10<sup>-3</sup> | F(1,17)=22.87<br>p<10<sup>-3</sup> |

The intra-personal variability of the responses was not affected by the posture for the haptic modality (α_upright_=6.2±6.1 mm, α_supine_=6.6±6.0 mm, posture effect: F_(1,17)_=0.12, p=0.73), but significantly increased in the supine position for the visual experiment (α_upright_=3.5±3.2 mm, α_supine_=4.8±4.7 mm, posture effect: F_(1,17)_=6.81, p=0.02).

In conclusion, in this experiment we found that patterns of distortion of both visual and haptic perception were invariant when expressed in an egocentric reference frame, but not when expressed in an exocentric one.

### Experiment 3: Gravity’s Effect on Visual and Haptic Perception

While the visual inputs are not different on ground and in weightlessness, the forces exerted against the virtual constraints during haptic exploration might be different in 0g due to biomechanical and neurophysiological effects. We therefore first analyze the changes in the contact forces between the subject’s hand and the virtual object and then the pattern of perceptive distortions (Figure 6A-C). The left plot of Figure 6A shows that vertical forces applied by the subjects on the upper and lower edge of the sensed object were modulated (F_(2,34)_=3.9, p=0.02) by the experimental conditions (1G, 0G, Supported). As expected, upward and downward forces increased and decreased respectively in microgravity (post-hoc 1G Vs 0G, p=0.02), coherent with a reduction of the weight of the upper limb. When the weight of the arm was supported (see methods), the vertical forces also tended to differ from 1G condition (post-hoc Supp Vs 1G p=0.09) and were modulated in the same direction as in 0G (post-hoc Supp Vs 0G, p=0.29). Horizontal forces were also significantly affected by the experimental condition (F_(2,34)_=6.32, p<0.01; Figure 6A, right plot), with a significant increase of the contact forces in microgravity with respect to the 1G and Support conditions.

**Figure 6:**
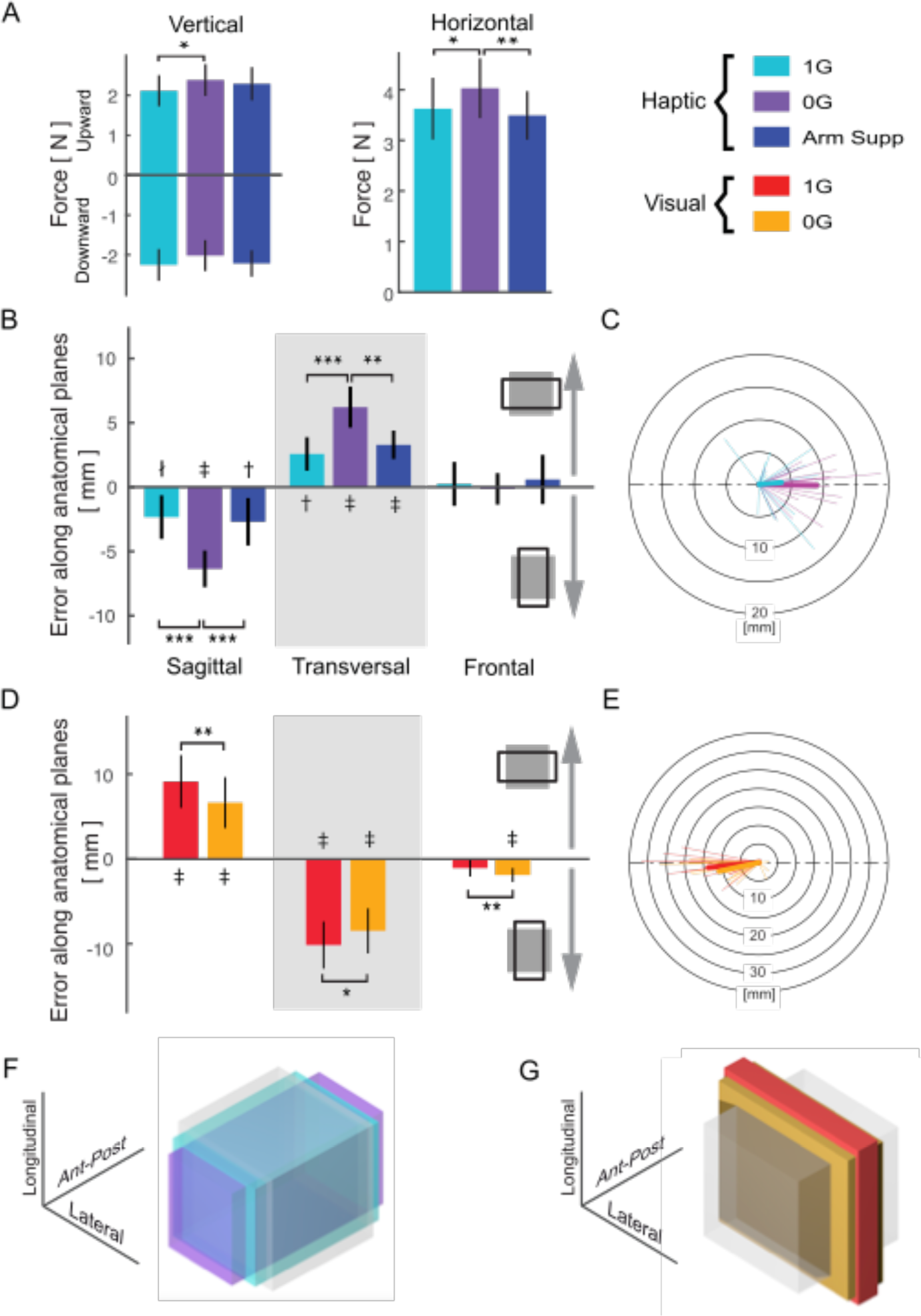
Results of the microgravity experiments for the haptic (A-C and F panels) and visual (D-E and G panels) tasks. A) Contact forces in the three experimental conditions: normogravity (1G), microgravity (0G) and with a mechanical support of the arm (Supp). Left: Vertical forces generated against the upper and lower edges of the rectangle. Right: Horizontal forces generated against all other edges of the rectangle. B) and D) Errors in the three task planes. C) and E) are circular vectorial plots representing individual responses in the Sagitt+Transv=Frontal errors space. Thicker vectors represent the average vectorial response. Dashed diameters indicate the Sagitt+Transv=0 axis. Within-subject variability of the responses when considering all three planes together. F-G) Illustration of the perceptive-distortion (experimental results scaled up by 5) of reference cube (gray) in normal gravity and in microgravity. * : p<0.05 and ** : p<10^−2^ in the ANOVA. ∤, † and ‡ : p<0.05, p<10^−2^ and p<10^−3^ for the t-test ascertaining difference from zero.

This increase of the contact force in 0G, similar to what was previously observed in haptic tests during parabolic flights (Mierau et al. 2008), could be the result of a specific strategy aimed at keeping muscular tension, and hence muscle spindle sensitivity, similar to normal gravity conditions. This strategy would avoid the decrease in proprioception precision previously observed in weightlessness for ‘open-chain’ motor tasks, for which the same strategy could not be adopted, resulting in a decrease in muscle tension (Clément and Reschke 2010). This hypothesis well matches the fact that the precision of haptic responses was not significantly affected by the experimental condition (response variability: 1G 6.8±2.6, 0G 7.1±3.1, Sup 6.4±2.9; F_(2,34)_=1.75, p=0.19), suggesting that neither microgravity nor the arm support significantly interfered with the subjects’ ability to perform the task. This lack of microgravity effect on haptic precision appears in line with the results of previous orbital experiments (McIntyre and Lipshits 2008).

Importantly, the results about the vertical contact forces and responses’ variability suggest that the ‘arm support’ condition successfully mimicked the expected lightening of the arm observed in microgravity. Therefore, if haptic perceptive distortions (constant errors) are affected by microgravity, but not by the arm support, they would not be directly ascribable to the biomechanical action of microgravity on the arm.

The comparison of the constant errors in the three experimental conditions, reported in Figure 6B, clearly shows that the perceptive distortion characterizing haptic perception in the Sagittal plane was significantly amplified (became more negative) by microgravity, but was not affected by the arm support (condition effect F_(2,34)_=12.49, p<10^−4^), suggesting a perceptive rather than biomechanical effect. Similarly, the haptic distortion in the Transversal plane was amplified (became more positive) in 0G and was not affected by the support, either (condition effect F_(2,34)_=11.13, p=<10^−3^). Finally, the lack of distortion in the Frontal plane persisted independent of the gravitational and support condition (F_(2,34)_=0.33, p=0.71). Figure 6C shows a clear increase of the global haptic distortion amplitude in 0G (bootstrap p<10^−4^) and very similar direction between the haptic distortions observed in the two gravitational conditions (bootstrap on the angle θ between the two vectors: p=0.64; R_0/1_=7.2).

For the visual tasks, Figure 6D shows that, as for the haptic sense, microgravity significantly modulated the perceptive distortions. More precisely, the large errors characterizing both sagittal and transversal planes in 1G were significantly reduced in weightlessness (F_(1,17)_=15.41, p=0.0011 and F_(1,17)_=7.87, p=0.012 respectively). In the frontal plane, a small but significant height underestimation appeared in 0G (F_(1,17)_=9.531, p=0.007). Figure 6E shows that the global visual distortion amplitude tends to decrease in microgravity, but not significantly (bootstrap p=0.15) and thatthe angle between the distortions in the two gravity conditions (6.7±5°) is not significantly larger than zero (bootstrap p=0.08): the probability of observing these data is eleven times larger under the hypothesis that the two vectors’ orientation representing the direction of the distortion are identical compared to the alternative hypothesis that they are different (R_0/1_=11.2). The analysis of the variable component of the errors shows that microgravity did not significantly affect subjects’ visual precision (F_(1,17)_=4.3, p=0.054), although the response variability tended to increase from 4.4±2.5 to 5.2±2.4 mm.

In neither haptic nor visual 0G tasks the amplitude of the distortions appears to change over the parabolas (trial number effect on haptic errors: Sagittal F_(4,60)_=0.79, p=0.54; Transversal F_(4,60)_=0.23, p=0.92; Frontal F_(4,60)_=0.49, p=0.74; and on visual errors Sagittal F_(4,68)_=1.23, p=0.30; Transversal F_(4,68)_=0.60, p=0.67; Frontal F_(4,68)_=0.63, p=0.64) suggesting a lack of significant adaptation to microgravity during the experiment duration.

The qualitative comparison of Figure 6F and Figure 6G shows that the effect of gravity on both sensory modalities mainly consists of a stretch of depth perception with respect to normo-gravity conditions (an increase in slenderness for haptic; a decrease in stubbiness for visual). The direct quantitative comparison of the effect of microgravity between the two groups of subjects of the visual and haptic experiments (Figure 7A) shows similar modulations of the perceptual distortion for both senses (Wilks’ Lambda=0.91, F_(3,32)_=0.96, p=0.42). Although the amplitude of the microgravity effect tends to be larger for haptic than for visual perception (bootstrap, p=0.06), the average directions of the microgravity effect on visual and haptic sense appear very similar (Figure 7B): the angle θ between the two vectors representing the average effect of gravity on the two modalities is only 15.6±15.6° and not significantly larger than zero (bootstrap, p=0.14). When considering the range of all possible θ (±180°), Bayesian statistics suggest that the observed data are 5.2 times more likely under the hypothesis that θ=0° (H_0_) than under the hypothesis θ≠0° (H_1_). As shown in Figure 7C, assuming that the gravity effect is identical for the haptic and visual modality (both in terms of direction and amplitude) predicts indeed distortions in microgravity indistinguishable from the observed results (Wilks’ Lambda=0.82, F_(4,14)_=0.77, p=0.56).

**Figure 7:**
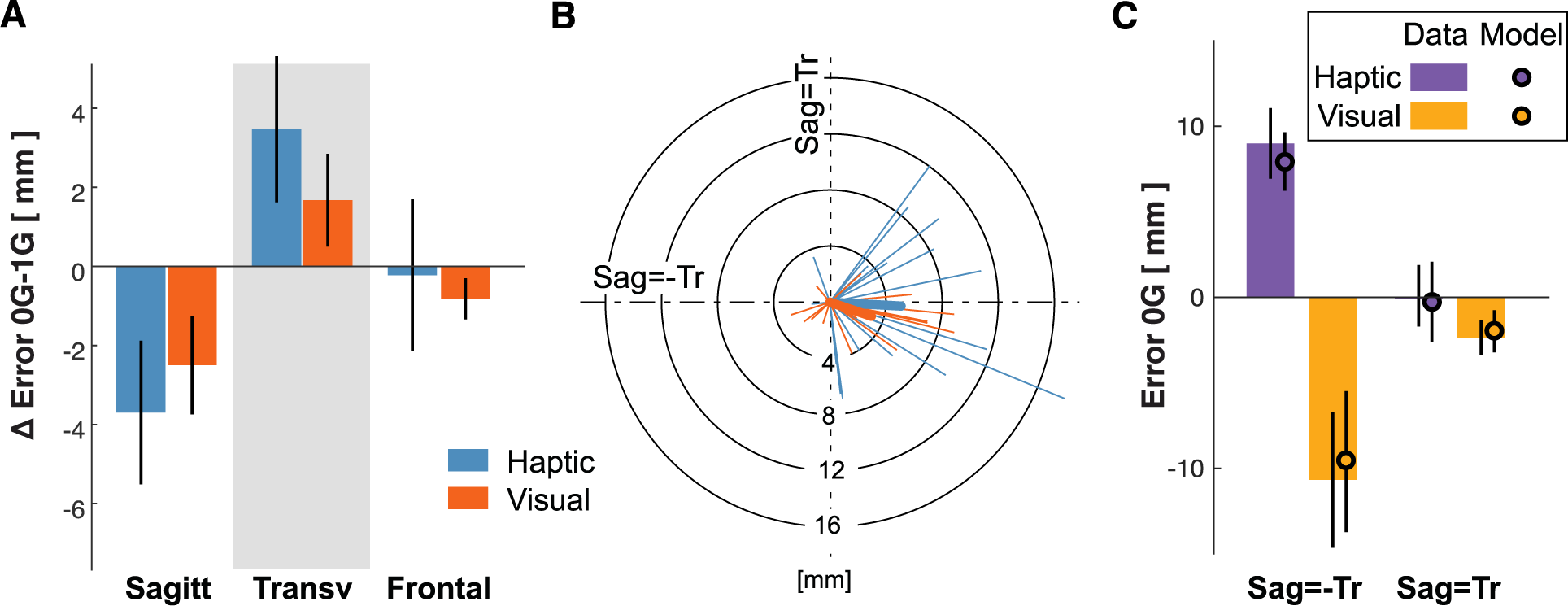
Comparison of the effect of microgravity on the Haptic and Visual senses. A) Difference between the constant errors made by the subjects in the 0G and 1G conditions for the tasks in the three anatomical planes. B) Vectorial representation of the gravity effect in the *Sag+Trans=Front* errors space. Thicker vectors represent the average response. Dashed line indicates the *Sag=-Trans* and *Sag=Trans* axes. C) Errors observed in 0G and predicted from 1G data assuming an identical effect of microgravity on both haptic and visual perception. The values correspond to the error projection on the *Sag=-Tr* and *Sag=Tr* axes of panel B. Vertical wiskers represent 95% confidence interval.

To summarize, the parabolic flight experiments show that, although very different perceptive distortions characterize vision and haptic sense in normal gravity conditions, the effects of microgravity on each of those distortions are in the same direction for the two sensory modalities.

### Results Summary

Experiment 1 revealed strong and opposite distortions between haptic and visual perception of 3D geometry. Subjects visually underestimated an object’s depth with respect to both height and width, whilst overestimating depth when exploring the object haptically. In Experiment 2 the comparison of seated versus supine body orientation clearly showed that both visual and haptic distortions align with the subject’s body rather than with gravity. Experiment 3, conducted during parabolic flight, showed a clear effect of microgravity on both haptic and visual distortion while preserving their respective ‘terrestrial’ direction. Importantly, the incremental effect of microgravity was along the same axis and in the same direction for visual and haptic tasks, despite the fact that the perceptive errors in normo-gravity were in diametrically opposite directions: weightlessness *increases* the haptic *over*-estimation of depth with respect to width and height and *decreases* the visual *under*-estimation of depth with respect to width and height.

## Discussion

The experiments presented here aimed to understand how gravity affects the perception of 3D shapes. We extend previous studies restricted to vision to include haptic sensation, by using the same experimental paradigm for the two modalities. Hereafter, we discuss our empirical results in normal gravity and in microgravity with respect to the literature and how they support the idea of a modality-independent role of gravity in interpreting incoming sensory signals.

### Haptic and Visual perception in normo-gravity conditions

Individually, the visual and haptic distortions observed here are consistent with previous findings obtained without using head-mounted displays or haptic devices, supporting the validity of the present experimental paradigms. Our haptic results concur with overestimation in the radial dimension observed for haptic tasks performed in the horizontal plane (Lipshits et al. 1994, Armstrong and Marks 1999, Fasse et al. 2000, Henriques and Soetching 2003). Similarly, the reported visual underestimation of depth corresponds to what has been previously observed for this modality in the horizontal plane (Wagner 1985, Loomis and Philbeck 1999). Surprisingly, we observed no significant distortion in the frontal plane, as one might have expected given the well-known horizontal-vertical, or “L”, illusion (Avery and Day 1969). The object being situated in front of the right shoulder, rather than straight ahead, might explain this discrepancy: in the latter position, vertical lines would more likely be associated with depth (Girgus and Coren 1975), than in the former, where horizontal lines may also contribute to depth perception.

Collectively, these results represent, to our knowledge, the first direct quantitative comparison of visual and haptic perception of 3D shapes. Although perceptual biases were known to differ between visually and haptically guided pointing (vanBeers et al 1999, Liu et al. 2018), no previous experiment has shown, as we do here, that the perceptual distortions are exactly in the opposite direction for the two senses.

Moreover, our second set of experiments on body orientation shows that visual and haptic distortions remain aligned to the subject anterior-posterior axis and opposite to each other when supine. Although in apparent contradiction with the effect of body tilt shown for a variety of visual tasks (Marendaz et al. 1993, Leone 1998, Barnett-Cowan et al. 2015), and with the effect of the direction of external forces in haptic tasks (Wydoodt et al. 2006), the present invariance of the ego-centric perceptive distortions is consistent with previously reported body-centered haptic perception (Gurfinkel et al. 1993, Dupin et al. 2018) and eye-centered encoding of shape/position of visual object (Averly and Day 1969, Howard et al. 1990, McIntyre et al. 1997, Henriques et al. 1998, Vetter et al. 1999). Furthermore, the present lack of effects of the body orientation on the egocentric visual and haptic 3D shapes perception is consistent with the previously reported absence of effects of tilting the body on unimodal visual and proprioceptive tasks, as opposed to cross-modal tasks (Bernard-Espina et al. 2022).

Our empirical observations of invariant, diametrically opposite visual and haptic egocentric distortions represent a novel and intriguing finding. It suggests that, in addition to causes associated with modality-specific mechanisms, such as eye vergence signals integration for vision (Murdison et al. 2019) and kinematics of the exploratory movements for haptic (Armstrong and Marks 1999, McFarland and Soechting 2007), the distortions characterizing the two modalities might have a common origin, probably linked to the continuous interaction between the information originating from the two sensory systems during everyday object manipulation: haptic distortions might compensate the perceptive visual bias, and viceversa, but this specific hypothesis remains to be tested.

### The role of gravity in 3D object perception

Although the ego-centric patterns observed for visual and haptic distortions would suggest that an external cue, such as gravity, should not influence shape perception, the strong microgravity effects observed in our parabolic-flights experiments clearly show that the contrary is, in fact, true. How can these apparently contradictory results be reconciled? We have shown that the observed effects of microgravity on both haptic and visual perceptive distortions are not directly ascribable to a decrease in their precision, or, specifically for the haptic task, to the mechanical action of gravity on the arm (arm support condition and supine haptic experiment). Moreover, the remarkable similarity between microgravity’s effects on visual and haptic distortions makes it unlikely that they might be caused by independent effects of gravity on the two sensory systems, as has been proposed for the results of previous microgravity experiments: modification of proprioceptive-tactile receptors’ output for haptic tasks (Lipshits et al. 1994) and deviation of the vertical position of the eye for visual tests (Clément et al. 2008, 2013, Bourrelly et al. 2015, 2016). A more parsimonious and likely explanation is that microgravity’s effect on the two senses has a common origin. This modality-independent phenomenon could be due to an effect of gravity on an internal representation of the egocentric 3D space used to interpret the object-related inputs, independent of the sensory system from which they originate. Clear evidence that internal models of space affect the interpretation of incoming sensory information in a Bayesian fashion has been extensively reported. Good examples include the ‘Ames room’ and the Müller-Lyer visual illusions based on prior knowledge about the geometry of constructed environments (O’Reilly et al. 2012) or the cutaneous Rabbit illusion (Goldreich et al. 2007). The idea that the same internal representation of space is used for both visual and haptic perception is also consistent with Müller-Lyer, inverted-T and L-shape illusions observed for both modalities (Taylor 2001, Gentaz and Hatwell 2004).

In conclusion, the present microgravity experiments testing visual and haptic shape perception with identical tasks provide the first direct experimental evidence of an effect of the gravitational signals on the modality-independent processing of spatial information, which could be only hypothesized in previous unimodal studies (Clement et al. 2009, 2012, 2013).

### Neural mechanisms

To understand how gravity affects this internal representation of space, it is important to consider the neural structures involved in 3D shapes perception. Brain imaging and electrophysiological studies have shown complex neural networks underlying object perception (Figure 8A): although 3D object images are known to activate the visual Lateral Occipital Complex, the same area is active during haptic shape recognition. Similarly, the areas commonly associated with haptic object perception, (primary and secondary somatosensory cortex, ventral premotor region, and Brodmann area 5) are activated also by images of manipulable objects. These cross-modal activations are mediated by the intraparietal sulcus (IPS) activity which is even enhanced during cross-modal, visuo-haptic object recognition (Grefkes et al. 2002). Electrophysiological studies have shown IPS activities consistent with recurrent neural networks able to perform cross-modal sensory transformations (Pouget et al. 2002, Avillac et al 2005), supporting its role in reconstructing a visual representation of a haptically sensed object and vice-versa (Figure 8B). The coexistence of a visual and a haptic object representation is consistent with the concurrent representations of reaching/grasping tasks aimed at statistically maximizing precision (McGuire and Sabes 2009, 2011, Tagliabue and McIntyre 2011-2014).

**Figure 8:**
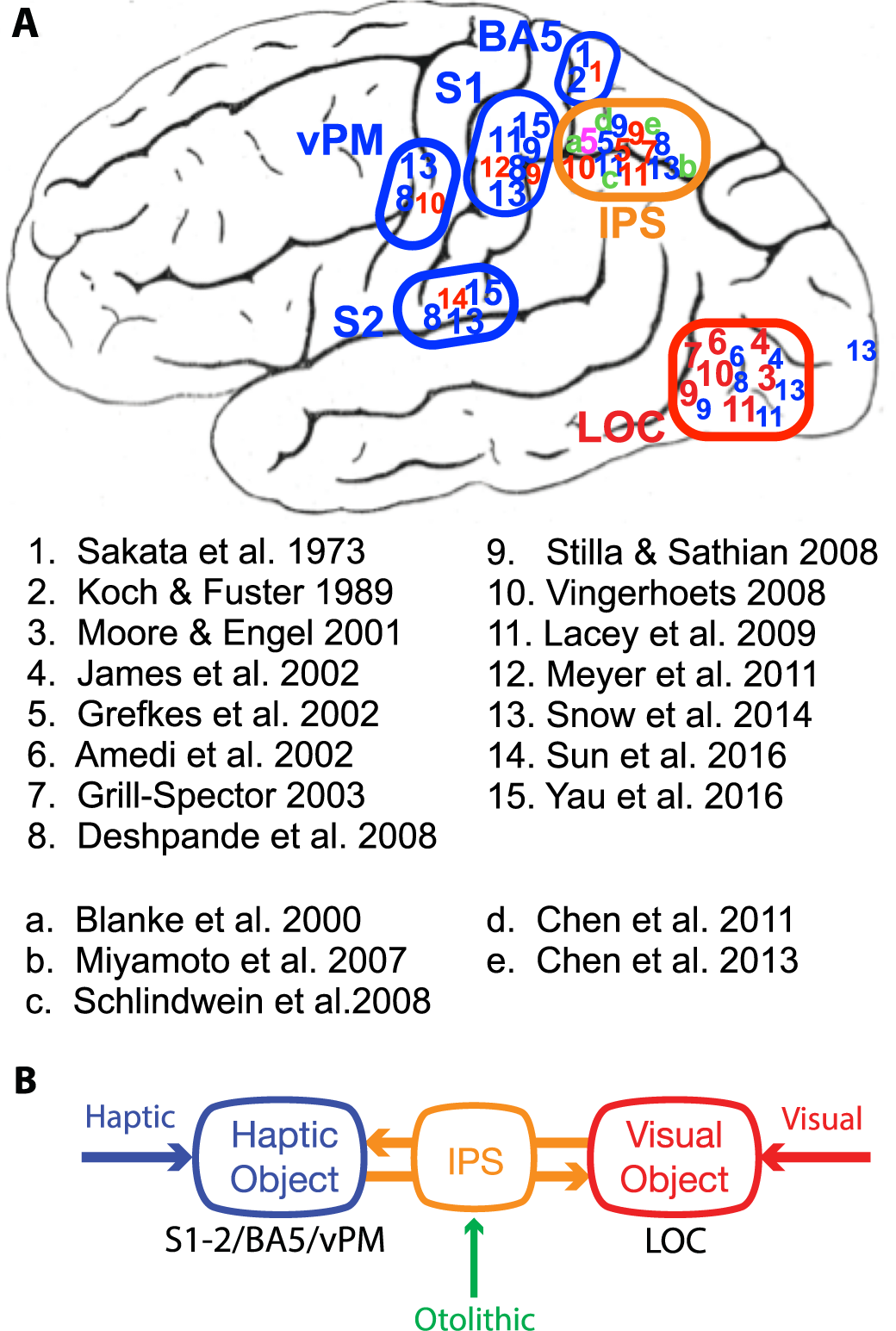
A) Brain cortical areas associated to haptic (blue), visual (red) and cross-modal (violet) objects’ perception. The regions primarily involved in haptic objects representation are the primary and secondary somatosensory areas (S1 and S2), the Brodmann area 5 (BA5), and the ventral premotor (vPM) area. The 3D object visual representation is known to reside in the lateral occipital complex (LOC). Numbers’ location represents the areas in which the corresponding study observed a neural activation during object perception tasks. Green letters represent studies reporting otolithic projection in the intraparietal sulcus (IPS) area. B) Simplified scheme of a putative neural network involved in objects perception. The IPS is connected to both haptic and visual object representation areas (S1/S2/BA5/vPM and LOC respectively). Its interaction with each of the two internal object representations would contribute to spatially interpret the haptic or visual sensory inputs during objects acquisition. IPS would also allow building a visual representation of the object from haptic signals and vice-versa. Otolithic signal affect IPS activity, thus indirectly both visual and haptic object representations.

To transform a visually-acquired object into a stable haptic representation (and vice-versa), despite potential independent movements of the two sensory systems, IPS must constantly integrate information about the eye-hand kinematic chain and the body position in space. In other words, the IPS contributes to build a stable internal representation of the egocentric space despite eye, head, and body movements (Andersen et al. 1997, Cohen and Andersen 2002, Land 2014).

The similar effect of gravity on both visual and haptic object perception observed here could then be explained by a role of gravitational/otolithic inputs in building the IPS internal representation of space. This idea is in line with the results of several electrophysiological studies in monkeys and humans reporting otolithic projections to IPS area (Figure 8) and with the observed head-tilt effect on the ability to perform visuo-proprioceptive transformations (Tagliabue and McIntyre 2011, 2013, Burns et al. 2011, Bernard-Espina et al. 2022).

### Conclusions

Our study provides a better understanding of human perception of 3D geometry. For the first time, we have shown a tight link between the axes of distortions characterizing visual and haptic object perception, even if the distortions are in opposite directions. This novel finding, together with the state-of-the-art knowledge about the neural circuitry underlying object perception with reciprocal activations of the visual and haptic systems, suggests a common origin for the observed perceptual biases. This idea is further strengthened by the observation that microgravity has the same incremental effect on visual and haptic object perception, consistent with the existence of a modality-independent perceptive mechanism: independently from the originating sensory system, the incoming object information would be treated by neural networks of the parietal cortex and interpreted through an internal representation of the egocentric 3D space, shaped by gravity (otolithic) signals. These microgravity experiments, therefore, provide fundamental cues to better understand the neurophysiology of perception “on Earth”. They suggest that unimodal 3D object perception might not actually exist, suggesting that restraining future investigations to the brain areas associated with a single sensory modality would be a clear limiting factor in understanding the neural mechanisms underlying 3D perception.

## Acknowledgment

The authors thank M. Patrice Jegouzo for his technical help in designing the experiments. This work was supported by the Centre National des Etudes Spatiales. This study contributes to the IdEx Université de Paris ANR-18-IDEX-0001.

## Appendix I: Graphically Representing Distortion

As we stated in the methods section, it is not possible to univocally quantify the absolute distortion for each of the three dimensions, but only with respect to the other two dimensions. However, in order to provide in the results section a graphical representation of the perceptive distortions, the following method was used to compute the dimensional errors. We first solved the system of equations of Table *1* reported below in the matrix form.

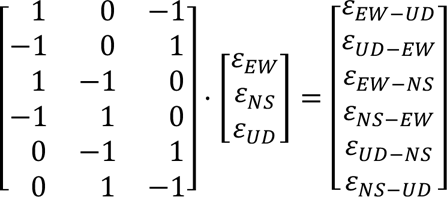

If we call *A* the matrix of linear coefficient, then the solutions of this underdetermined problem are

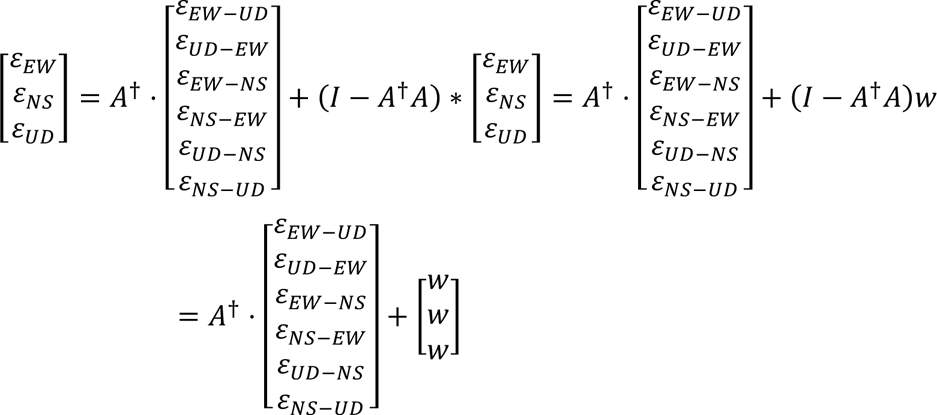

Where the *A*^3^ is the pseudo inverse of *A* and *w* is a free scalar parameter that reflects the fact that the observed results can be explained by an infinity of triplets of dimensional distortions differing by isotropic component, *w*, only (underdetermination of the problem).

To define a set of dimensional distortions, (ε*_EW_*, ε*_NS_*, ε*_UD_*) to be used for a graphical representation, we arbitrary decided to select the solution that minimizes the Euclidean norm of the distortion vectors.

Although the *w* parameter cannot be univocally defined, the difference between the errors along the three dimensions are correctly quantified and then used to test the anisotropy of the perceptive errors. The dimensional errors, however, cannot be rigorously compared between postures or gravitational conditions, because the differences between experimental conditions could be due to differences in defining the *w* parameter for each condition.

